# The alkane 1-monooxigenase gene *alkB* of *Pseudomonas* sp. FF2 is upregulated during colonisation of *Arabidopsis thaliana* leaves

**DOI:** 10.1101/2025.03.25.645160

**Authors:** Rudolf O. Schlechter, Laura Voß, Evan J. Kear, Mila Oeltjen, Mitja NP. Remus-Emsermann

## Abstract

Bacteria on leaf surfaces encounter highly variable access to nutrients and water. This oligotrophic environment is partly due to the presence of cuticular waxes that render the leaf surface hydrophobic, reducing the leaching of nutrients and water loss from inside the leaf. Bacteria have evolved adaptations to survive under these conditions. While *alkB*, an alkane hydroxylase gene, is widely prevalent in leaf-associated bacteria, its role and activity is unclear. Here, we developed a bioreporter in *Pseudomonas* sp. FF2 (PFF2) to investigate *alkB* promoter activity in diesel and on *Arabidopsis thaliana* leaves. In general, *alkB* promoter activity is highly heterogeneous, with a subpopulation exhibiting strong activation, suggesting bet-hedging in alkane metabolism. *In planta*, the promoter remained active over the course of seven days, indicating constant access of alkanes over time. Single-cell fluorescence intensity was heterogeneous, reflecting differences in microhabitats on the leaf surface or bet-hedging. While our results support a role of *alkB* in bacterial adaptation to the phyllosphere, direct evidence of cuticular wax degradation is missing. Future studies should trace the incorporation of plant-derived aliphatic compounds to elucidate the potential use of alkanes and other aliphatic compounds as resources for bacteria in the leaf environment.

## Introduction

The phyllosphere represents the microbial habitat that is formed by the surfaces of all aboveground plant organs, such as stems, flowers, fruits and leaves (Sohrabi et al. 2023). Leaves are the most prevalent part of the phyllosphere and usually, the term phyllosphere is used synonymously with the leaf surface habitat (Leveau 2019). On the macroscale, factors such as UV radiation, fluctuating temperatures, and variable weather conditions make the leaf surface an extreme environment to colonise. On the microscale, challenges include variable nutrient and water availability (Schlechter, Miebach, and Remus-Emsermann 2019), largely influenced by the diverse and highly hydrophobic structure of plant surfaces (Yeats and Rose 2013; Schreiber 2010). Despite these challenges, the phyllosphere is home to many microorganisms, predominantly bacteria, which colonise this environment at densities of 10^6^ and 10^7^ cells per cm^2^ of leaf surface (Schlechter, Miebach, and Remus-Emsermann 2019). Bacteria in the phyllosphere have evolved various adaptation strategies to withstand these adverse conditions. Examples include UV protection via pigmentation and DNA repair mechanisms (Schlechter, Miebach, and Remus-Emsermann 2019), motility and biosurfactant production (Kunzler et al. 2024) and highly specific resource uptake mechanisms (Delmotte et al. 2009). As nutrients are a limiting growth factor on the leaf surface, specialised mechanisms for nutrient acquisition are critical and a fitness advantage. For example, *Sphingomonas* express TonB receptors to increase nutrient uptake on the leaf surface, while other taxa such as the facultative methylotrophic *Methylobacterium* spp. can rely on the use of methanol as a byproduct of the plant cell wall biosynthesis (Yurimoto, Shiraishi, and Sakai 2021; Delmotte et al. 2009). Although the phyllosphere is usually considered an oligotrophic environment with low amounts of photosynthates leaching to the leaf surface (Remus-Emsermann et al. 2011; Tukey and Mecklenburg 1964), it is rich in alkanes (Schreiber and Schönherr 2009). These alkanes are components of the leaf cuticle, the uppermost layer of the leaf which serves as a barrier to reduce water and nutrient loss, and protect against biotic stress (Yeats and Rose 2013; Zeisler-Diehl, Barthlott, and Schreiber 2020).

The plant leaf cuticle is primarily composed of cutin, a biopolyester composed of hydroxy- and hydroxyl epoxy-fatty acids, and waxes that include aliphatic compounds, with a highly diverse mixture of carbon chains ranging from C16 to C32 (Schreiber and Schönherr 2009). Many bacteria colonising the phyllosphere have been found to be able to degrade hydrocarbons (Simisola Oso et al. 2019). Mostly, this trait is conferred by the *alkBFGHIJKL* and *alkST* operons (Nieder and Shapiro 1975; Grund et al. 1975). The *alkBFGHIJKL* operon is controlled by the *alkB* promoter (P*alkB*) and the *alkST* is under the control of the transcription factor AlkS. AlkS activates P*alkB* in the presence of alkanes. The activity of P*alkB* has previously been found to be dependent on the availability of alkanes in the growth medium and is therefore well suited as a proxy for the activity of *alkB* (Yuste et al. 2000). The *alkB* gene is one of the most ubiquitous genes involved in alkane degradation and is widely distributed in many phylogenetically diverse bacteria. Moreover, *alkB* is often used as a representative of the entire alkane degradation pathway and has been used to assess the composition of alkane-degrading communities, as well as to determine their response to hydrocarbon inputs (Smith et al., 2013). The *alkB* gene codes for a membrane-bound non-heme diiron alkane 1-monooxygenase, which is involved in the first step of an oxidation pathway to transform alkanes into fatty acids (Guo et al. 2023). Given the ubiquity of *alkB* among phyllosphere-associated bacteria, it is hypothesised that AlkB-mediated alkane degradation offers a fitness advantage for bacteria in this environment (Gandolfi et al. 2017).

Previous studies have shown that certain phyllosphere-associated bacteria, including several *Pseudomonas* spp., can grow on diesel, which is chemically similar to the hydrocarbons found in the cuticle (S. Oso et al. 2021; Simisola Oso et al. 2019). One of these Pseudomonads, *Pseudomonas* sp. FF2 (PFF2), isolated from romaine lettuce (Burch et al. 2016), has been shown to utilise diesel as its sole carbon source and harbours an *alkB* gene (S. Oso et al. 2021). PFF2 is part of the *P. fluorescens*-lineage and subgroup and is closely related to *P. extremaustralis*. In this study, we hypothesised that PFF2 expresses *alkB* on plant leaves to degrade cuticular alkanes. To that end, a bioreporter was developed by constructing a plasmid containing a green fluorescent protein (GFP) gene under the native *alkB* promoter of PFF2 to monitor the *alkB* gene activity during leaf surface colonisation at the single-cell resolution. The functionality of the bioreporter was tested *in vitro* and on *Arabidopsis thaliana* leaves. The fluorescence of single cells was analysed to provide insights into the activity of the *alkB* promoter and as a proxy for the activation of bacterial *alkB*-mediated alkane degradation in the phyllosphere.

### Experimental Procedures

#### Strains, culture media and growth condition

Strains were routinely grown in lysogeny broth (LB (Luria/Miller), Carl Roth) or lysogeny broth agar (LBA (Luria/Miller), Carl Roth). Unless stated otherwise, *Escherichia coli* was grown at 37°C and was used for cloning and as donor for conjugation experiments (ST18), while the focal strain *Pseudomonas* sp. FF2 (PFF2) was grown at 30°C. Wherever appropriate, 50 µg mL^-1^ kanamycin (Km, Carl Roth) was used in selective media.

Bushnell Haas Broth (BHB, 409.6 mg/L MgSO_4_·7H_2_O, 26.5 mg/L CaCl_2_·2H_2_O, 1.0 g/L KH_2_PO_4_, 1.0 g/L K_2_HPO_4_, 1.0 g/L NH_4_NO_3_, 83.3 mg/L FeCl_3_·6H_2_O, pH 7.0) was used as a base medium for *in vitro* growth in diesel. BHB was supplemented with 1 % v/v diesel (commercial diesel, locally sourced). A chemical analysis of this diesel was not performed; therefore, its composition was not determined.

#### Alkane bioreporter construction and conjugation into PFF2

To monitor *alkB* promoter activity, we constructed the plasmid pFru97-PalkB-mClover3-[AAV] (pPalkB-GFP*), which expresses a destabilised version of the green fluorescent protein mClover3 under the control of a 300 bp native *alkB* promoter fragment from *Pseudomonas* sp. FF2 (PFF2). The reporter cassette was assembled via isothermal isothermal assembly and inserted into the broad-host-range vector pFru97 (Supplemental Methods). The assembled pPalkB-GFP* plasmid was then transformed into NEB Turbo competent *E. coli* cells (New England Biolabs) according to the manufacturer’s recommendations and were selected on LBA supplemented with 50 µg/ml Km. The plasmid was verified by Sanger sequencing (GENEWIZ Germany GmbH) and then transformed into chemically-competent *E. coli* ST18 before plating on LBA supplemented with Km + 5-aminolevulinic acid (5-ALA, 50 mg/ml) to select for transformants (Thoma and Schobert 2009)

Following transformation into *E. coli* ST18, the pPalkB-GFP* plasmid was transferred into PFF2 via biparental mating (Supplemental Methods) (Schlechter *et al*. 2019). Transconjugants were selected on kanamycin, counter-selected against *E. coli*, and screened by PCR for plasmid presence and absence of donor contamination. The resulting strain, PFF2_PalkB-GFP*_, was used for all subsequent experiments.

#### Diesel utilisation assay and P*alkB* activity in Bushnell-Haas broth and LB

The newly developed alkane bioreporter was first tested *in vitro*. To that end, 5-mL overnight cultures of PFF2_P*alkB*-GFP*_ grown in LB + Km at 30°C were used to inoculate BHB medium supplemented with 1% v/v diesel (S. Oso et al. 2021). The overnight cultures were washed by centrifugation for 10 min at 5,000 × *g* and then resuspended in 5 ml BHB (*N* = 3). PFF2_P*alkB*-GFP*_ was grown in 50 ml BHB with or without 1% v/v diesel. BHB + 1% v/v diesel without bacteria was used as a negative control. All cultures were supplemented with Km and incubated at 30°C with constant shaking at 300 r.p.m. Optical density at 600 nm was measured using a spectrophotometer (GeneQuant 100, Biochrom) in three-to-four-day intervals over a total time of 28 days. At days 1, 6, and 28, aliquots were samples and prepared for single-cell analysis. Furthermore, the fluorescence intensity of PFF2_P*alkB*-GFP*_ was measured in LB.

#### Growth of PFF2_P*alkB-GFP\**_ *in planta* and activity of P*alkB*

To determine *alkB* promoter activity during colonisation of *A. thaliana* leaves, PFF2_P*alkB*-GFP*_ was inoculated onto axenically grown *A. thaliana* (Supplemental Methods). *A. thaliana* Col-0 seeds were surface-sterilised and stratified at 4°C in the dark for two days. Afterwards, the seeds were sown onto ½ strength Murashige and Skoog (MS with vitamins, pH 5.8, Duchefa) 1.0% w/v plant agar (Duchefa) plates with each seed placed on a trimmed sterile pipette tip filled with solid MS agar. Seeds were grown in a plant growth cabinet (poly klima GmbH) at 21 °C in an 11/13 h photoperiod, with a light intensity of ∼120 μE m^-2^ s^-1^, and a 80% relative humidity. Eight days after sowing, seedlings were transferred aseptically into autoclaved Magenta GA-7 (Merck) plant tissue culture boxes containing 90 g of zeolite (Zeolith-100, Steinlando) and 45 ml of sterile¾ MS, with four seedlings per box.

Four-week-old plants were inoculated with an exponentially grown PFF2_PalkB-GFP*_ culture set at OD_600nm_= 0.05 in PBS using an airbrush sprayer (Ultra airbrush, Harder & Steenbeck GmbH & Co. KG; compressor, Sparmax TC-620X airbrush compressor). The inoculated boxes were then placed into the plant growth cabinet under the same controlled conditions.

Whole *A. thaliana* rosettes were sampled on day zero, two, and seven after inoculation, with six to eight biological replicates sampled each time. PBS were added to the samples, vortexed for 10 sec, sonicated for 5 minutes at 75% intensity in a sonication bath (Emmi-12HC, EMAG AG), and then vortexed for an additional 15 sec. An aliquot of the leaf wash was used to prepare a tenfold dilution series and to determine colony forming units (CFU) on LBA + Km. Plant fresh plant weight was recorded for normalisation of colony counts. The remaining suspension was used for microscopy.

#### Microscopy

Samples from *in vitro* and *in planta* experiments were prepared for microscopy by harvesting bacterial suspensions via centrifugation at 15,000 × *g* for 10 min at 4 °C. The supernatant was discarded, and the resulting pellet was resuspended in 50 μl of 4% paraformaldehyde solution (PFA, 4% w/v PFA in 1×PBS w/v, pH 7.2) and incubated for 1 hour at room temperature or overnight at 4°C. To remove residual PFA, the cells were washed three times by centrifugation at 15,000 × *g* for 5 min at 4 °C and resuspended each time in 1×PBS. After the final wash, the pellet was resuspended in 50 μl 1×PBS. Subsequently, 50 μl ethanol was added to the samples and the samples were analysed using microscopy within 24 hours after harvesting. Microscopy was performed on a Axio Imager.Z2 (Carl Zeiss) using an EC Plan-Neofluar 100 × /1.3 oil Ph3 objective using phase contrast and the Zeiss Filter 38 HE (BP 470/ 40, FT 495, BP 525/ 50), to acquire phase contrast images and mClover3 signals, respectively. Micrographs were acquired using a Axiocam 712 mono camera (Carl Zeiss) and the programme ZEN Blue 3.3 (Carl Zeiss).

#### Image processing

Quantitative image analysis was performed using custom macros to automate image processing in FIJI v. 2.3.0 (Schindelin et al. 2012). Raw images were imported and converted to 8-bit grayscale. A training data set was generated from random images for cell classification. To that end, phase contrast images were used for segmentation with a threshold applied based on the image type (*in vitro* or *in planta* images), set between one and four standard deviations below the mean pixel intensity. Masks were created and refined using morphological operations such as watershed segmentation, erosion, and dilation. Particles between 0.25 and 6.0 µm^2^ were detected, and each particle was manually classified as a bacterial cell or non-cell. The data was then used in machine learning model training.

For automated cell segmentation, images were converted to 8-bit grayscale, and phase contrast and green fluorescent channels were extracted. A mask was generated from the phase contrast as explained above. Particles were selected and mapped onto the green fluorescent channel for fluorescence intensity measurement. Additional measurements were recorded for cell classification using machine learning, including area, mean pixel intensity, perimeter, shape descriptors (major and minor axis, angle), circularity, Feret’s diameter (Feret, FeretX, FeretY, Feret Angle, MinFeret), aspect ratio, roundness, and solidity.

Background fluorescence was estimated from images that were thresholded using the percentile method. The resulting binary mask was inverted to exclude regions with bacterial biomass. Then, 50 randomly positioned 2×2-pixel squares were selected across the background regions, and the mean fluorescence intensity in the green fluorescent channels was measured. These values were used for downstream fluorescence correction.

#### Machine learning model development

Cell classification was performed using machine learning, with models trained on ground truth-labeled datasets. Using a custom macro script in FIJI/ImageJ, particles from a subset of images were manually classified as cells or non-cells, and particle descriptors were recorded to identify the most relevant predictors for classifying real datasets. Two machine learning models for each of the *in vitro* and *in planta* data set were trained to classify single cells: a logistic regression (LR) with L1 regularisation, and a random forest (RF) model. These models were developed using the R package *tidymodels (Kuhn and Wickham 2020)*. The LR model was implemented using the *glmnet* engine with an L1 penalty (Lasso regression), with the penalty parameter optimised over a logarithmic spaced grid from 10^-4^ to 10^-1^. Best penalty was then used to improve the model based on the higher area under the receiver operating characteristic (ROC) curve. The RF model was implemented using the *ranger* engine with 500 trees. The number of predictors sampled at each split and the minimum number of observations required in a node were tuned over 25 combinations, selecting the best parameters based on ROC performance. Single-cell data was split into two datasets: training (75%) and test (25%). Both models were trained using ten-fold cross-validation on the training data set, and the final models were evaluated on the test dataset. Performance metrics were estimated, including accuracy, precision, recall, F1-score, kappa statistics, area under the ROC curve (ROC-AUC), sensitivity, and specificity. Feature importance was assessed using variable importance plots to select the best models.

#### Single-cell fluorescence analysis

Fluorescence of single cells was analysed in R. Cells classified as positive by the best-performing machine learning model were retained for further analysis. Fluorescence values were corrected by subtracting the median background intensity from each cell’s fluorescence signal per image, and replicates with insufficient cell counts were excluded.

The distribution of fluorescence values was assessed using normal probability plots, comparing the observed fluorescence data with an expected normal distribution. Deviations from normality were evaluated using skewness and kurtosis, calculated with the *moments* package (Komsta and Novomestky 2022). A normal distribution is characterised by a skewness = 0 and kurtosis = 3. Positive skewness indicates a right-tailed distribution, while kurtosis > 3 suggests the presence of heavy tails (extreme values).

To quantify high-fluorescent cells and determine fold-change relative to a control, a baseline fluorescence threshold was calculated using the interquartile range criterion, defined as I = *Q*_75_ + 1.5 × IQR, where *Q*_75_ was the 75th percentile of fluorescence values from BHB-grown cells without diesel (*in vitro*), or time zero (*in planta*).

Statistical analyses were performed using the *rstatix* package (Kassambara 2023) at α= 0.05, and included normality tests (Shapiro-Wilk), homoscedasticity tests (Levene’s), and comparisons between fluorescence distributions using t-tests, ANOVA, and Tukey’s HSD. Kruskal-Wallis and Dunn’s tests were used as non-parametric alternatives. Results were visualised in cumulative distribution and probability plots, as well as comparisons of the highest 1% highest fluorescence values across time points.

## Results

### PFF2 growth *alkB* promoter activity *in vitro*

To investigate the growth of PFF2_P*alkB*-GFP*_ and the *alkB* promoter activity in diesel-supplemented minimal media, we cultured PFF2_P*alkB*-GFP*_ in BHB with 1% v/v diesel and in BHB without an added carbon source. PFF2_P*alkB*-GFP*_ reached a mean maximal OD_600nm_ of 0.61 ± 0.052 after 18 days (**Fig 1**). In contrast, growth was negligible in BHB alone, with no change in OD_600nm_.

**Figure 1.**
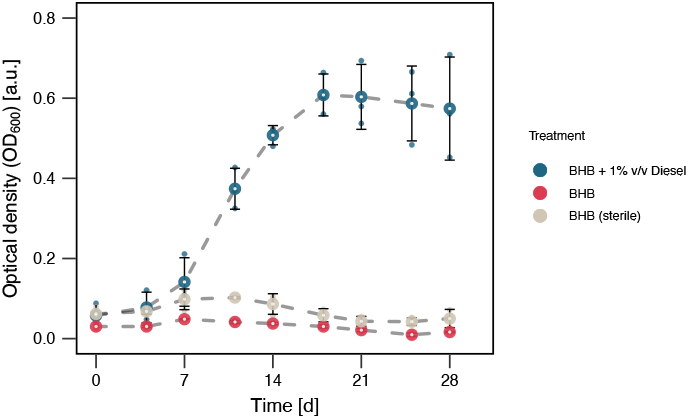
Growth of PFF2_PalkB-GFP*_ in diesel-supplemented media. Optical density (OD_600_) measurements of PFF2_PalkB-GFP*_ grown in Bushnell-Haas broth (BHB) supplemented with 1% v/v diesel, compared to BHB without a carbon source and a non-inoculated sterile control. Growth was monitored over time to assess the impact of diesel as a sole carbon source.

The increase in OD_600_ correlated with elevated *alkB* promoter activity, as measured by single-cell fluorescence imaging. A trained random forest model classified individual cells with high accuracy, precision, and ROC AUC (>90%), outperforming a logistic regression model marginally (**Fig S2A**). The most important classification features were the growth condition (BHB no diesel), particle solidity (area/convex area ratio), and aspect ratio (**Fig S2B and C**). Fluorescence distributions of PFF2_P*alkB*-GFP*_ were consistently right-skewed across all conditions (**Fig S3**). In diesel-grown PFF2_P*alkB*-GFP*_, skewness and kurtosis averaged 2.7 and 25.2, respectively, compared to 2.1 and 8.1 in non-diesel supplemented media. This strong positive skewness (>2) and kurtosis (>3) reflected a large proportion of low-fluorescence cells.

Over time, 46–90% of diesel-grown PFF2_P*alkB*-GFP*_ cells showed fluorescence values below 10% of the maximum measurable intensity (25.6 a.u., on a 256-pixel scale), compared to 66-83% in cells grown in BHB without diesel. As a comparison, 90% of cells grown in LB displayed low fluorescence values; however, maximal values were comparable to cells grown in BHB without diesel. This consistent pattern across conditions suggests that heterogeneity in *alkB* promoter activity is intrinsic to a population, regardless of growth condition.

To further assess fluorescence dynamics, we normalised fluorescence in diesel-grown cells to the fluorescence of cells grown in BHB alone medium at each time point. Median relative fluorescence remained low across time points (0.23–0.46-fold), but skewness and kurtosis increased especially at day 28 **(Fig 2A, Table 1)**. However, at one-day post inoculation, 19% of cells exceeded baseline fluorescence, with some cells reaching up to 3.6-fold higher fluorescence. By day six, only 3.9% of cells exhibited higher fluorescence, with a maximum of 2.4-fold above baseline. By day 28, relative fluorescence increased, with 11% of cells exceeding baseline fluorescence by up to 5.8-fold.

**Table 1.** Summary of *in vitro* single cell microscopy of PFF2 grown in BHB + 1% diesel.

| Time [dpi] | Biological Replicates | N cells | Median Rel. Fluorescence | IQR Rel. Fluorescence | Proportion High Fluorescence [%] | Max Rel. Fluorescence | Skewness | Kurtosis |
| --- | --- | --- | --- | --- | --- | --- | --- | --- |
| 1 | 3 | 3,839 | 0.46 | 0.122 | 18.5 | 3.6 | 1.44 | 5.93 |
| 6 | 3 | 5,721 | 0.23 | 0.055 | 4.6 | 2.4 | 1.46 | 5.83 |
| 28 | 3 | 10,988 | 0.23 | 0.067 | 3.78 | 5.8 | 3.30 | 24.5 |

**Figure 2.**
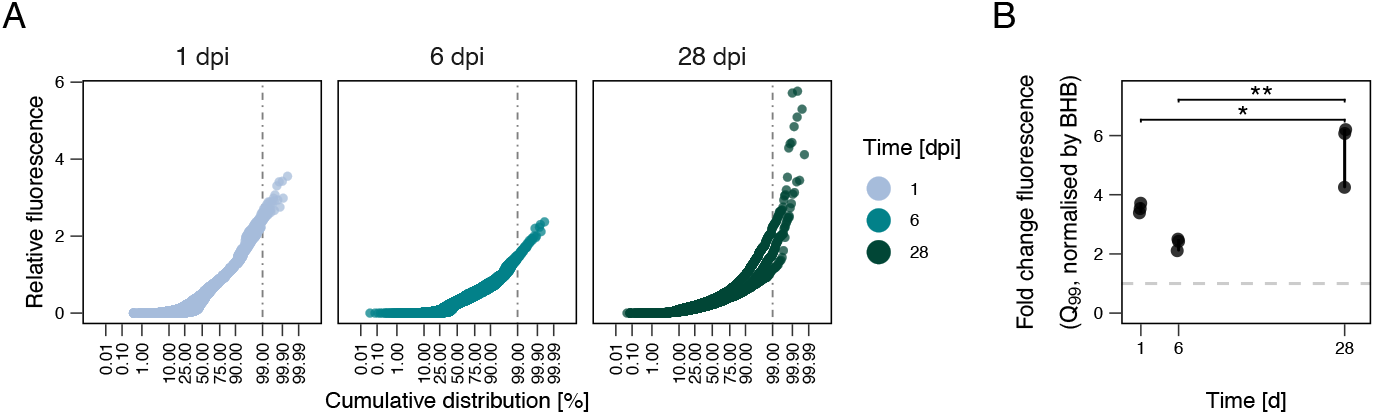
Relative fluorescence distribution and high-intensity cell subpopulations of PFF2_PalkB-GFP*_ in diesel-supplemented media over time. (A) Normal probability plot of PFF2_PalkB-GFP*_ single-cell green fluorescence in BHB + 1% diesel at 1, 6, and 28 days, relative to baseline fluorescence. Baseline fluorescence was defined using the interquartile range (IQR) criterion (*I* = Q3 + 1.5 × IQR) for cell populations incubated in BHB without diesel at each time point. The vertical line marks the 99th percentile of the fluorescence distribution. (B) Fold change in fluorescence of the top 1% of cells (99th percentile), normalised to baseline fluorescence. The horizontal line indicates a fold change of 1. Asterisks (*) denote statistically significant differences among groups (*p* < 0.05, ANOVA with Tukey’s post hoc test).

To determine whether highly fluorescent cells differed significantly from the overall populations, we analysed the top 1% of fluorescent cells across growth conditions **(Fig 2B)**. The median relative fluorescence of these cell subpopulations was higher than baseline at every sampling point (one-sample t-test, *p* < 0.05). Differences between time points were observed (ANOVA, *F*(2,6) = 18.43, *p* = 0.003), with cell fluorescence at day one and six being statistically lower than at day 28 (Tukey’s HSD, *p* < 0.05).

Taken together, these results indicate that while diesel supplementation supports growth of PFF2, *alkB* promoter activity is highly heterogeneous within the population. A small fraction of cells exhibited elevated fluorescence, particularly at early and late time points, while fluorescence was low in most cells. Thus, only a subset of cells actively engages in alkane degradation at a given time.

### Growth and promoter activity on arabidopsis

We assessed the dynamics of population growth and single-cell fluorescence over time in in *A. thaliana* to evaluate the heterogeneity of PFF2_P*alkB*-GFP*_ in the phyllosphere. Two independent experiments showed a consistent pattern of population growth **(Fig 3)**. At time zero, median CFU values ranged from 2.07–3.30 × 10^5^ CFU per gram of leaf (gFW). By day two, CFU increased 6- to 20-fold, reaching 1.40–4.15 × 10^6^ CFU gFW^-1^, and maintaining high levels by day seven, with a median of 3.38–6.35 × 10^6^ CFU gFW^-1^.

**Figure 3.**
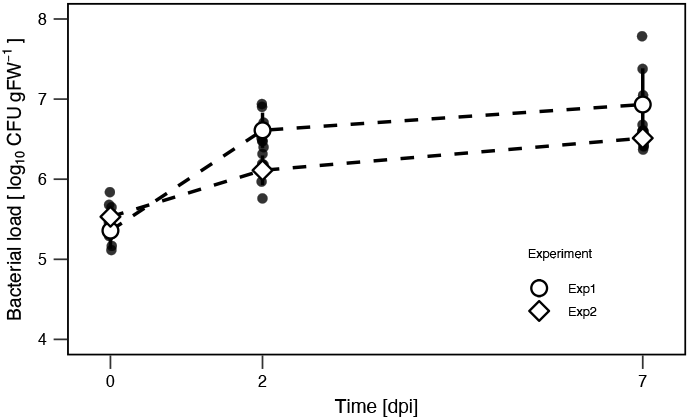
Population growth of PFF2P_alkB-GFP*_ on *A. thaliana*. Bacterial loads were quantified as CFU per gram of fresh weight across two independent experiments. (N = 8 biological replicates per experiment).

In parallel, we sampled bacterial cells and assessed their fluorescence as a proxy for *alkB* promoter activity using fluorescence microscopy. Similar to the *in vitro* dataset, a random forest model slightly outperformed a logistic regression model and was therefore used for cell classification **(Fig S4A)**. The most important classification features included particle solidity, aspect ratio, and roundness (**Fig S4B**,**C**).

Single-cell data showed a consistent pattern of increasing fluorescence heterogeneity over time across the two independent experiments. Fluorescence distributions became more right-skewed and with high kurtosis **(Fig S5)**. To evaluate changes in *alkB* promoter activity, we normalised the fluorescence values to the fluorescence of cells at time zero **(Fig 4A)**. At time of inoculation, cell fluorescence showed the lowest skewness and kurtosis **(Table 2)**, indicating a near-normal distribution. By day two, distribution of single-cell fluorescence became more asymmetrical, with increased skewness and kurtosis **(Table 2)**. This pattern persisted on day seven, revealing consistent heterogeneous PalkB activity. Within the cell populations, the proportion of highly fluorescent cells increased over time, with 8.2–36.1% of cells exceeding baseline fluorescence by day two and 16.9–36.2% by day seven.

**Table 2.**
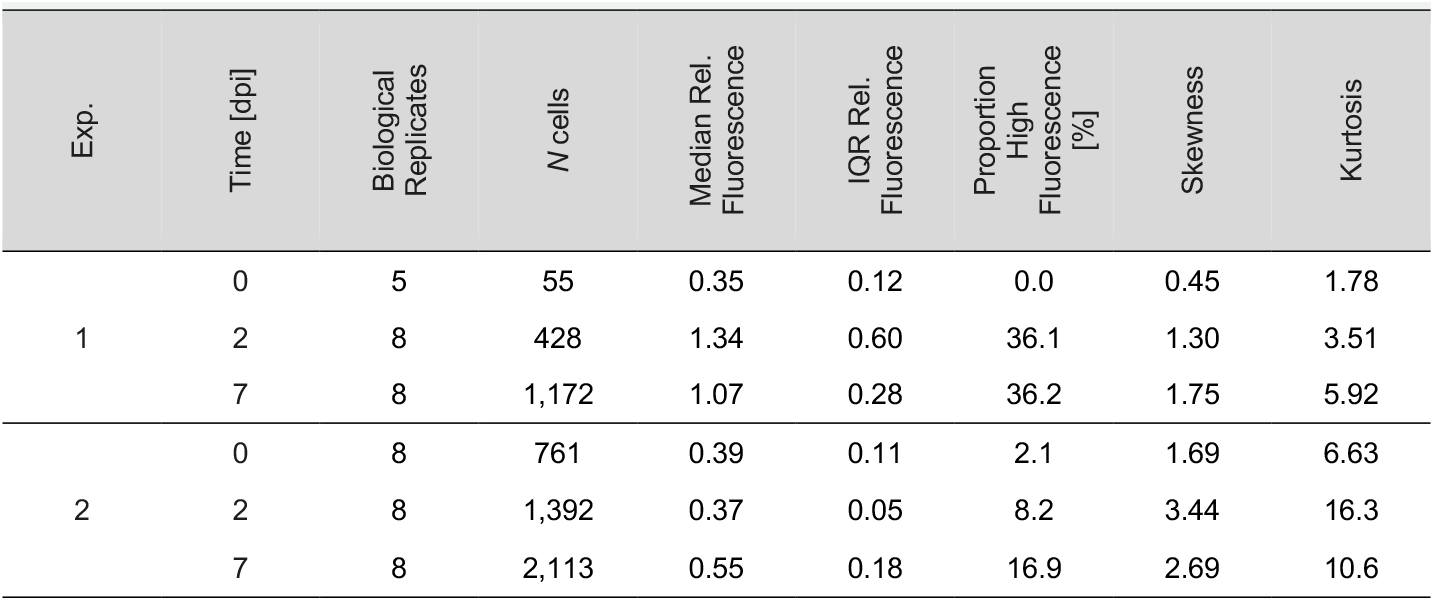
Summary of *in planta* single-cell microscopy.

**Figure 4.**
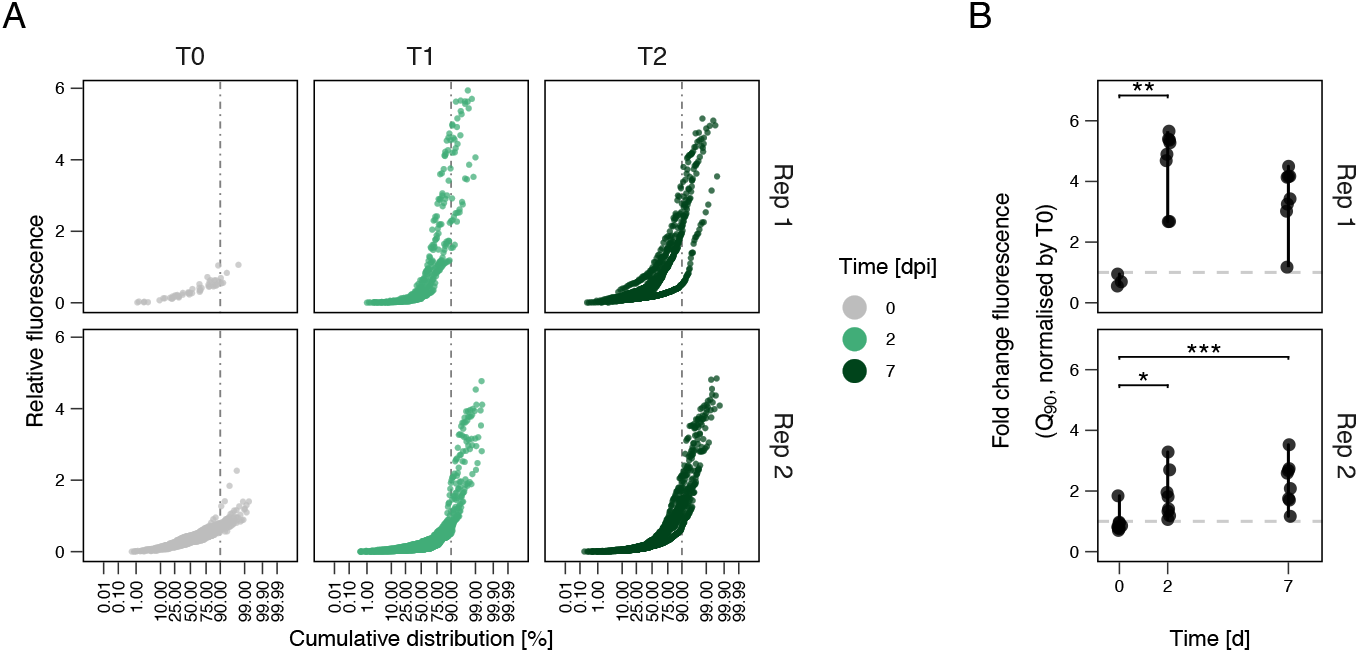
Distribution of relative single-cell fluorescence and *alkB* promoter activity in PFF2_PalkB-GFP*_ in *A. thaliana*. (A) Normal probability plot of PFF2_PalkB-GFP*_ single-cell GFP fluorescence at different sampling points (0-, 2-, and 7-days post-inoculation) relative to the baseline fluorescence. Baseline fluorescence was defined using the IQR criterion (*I* = q.75 + 1.5 × IQR) from cell populations at time zero (0 dpi). The vertical line indicates the fluorescence threshold below which 90% of the cell population falls. (B) Fold change in fluorescence of the top 10% most fluorescent cells (90% quantile), normalised to the baseline fluorescence. The horizontal line represents a fold change of 1. Asterisks indicate statistically significant differences among groups (* = p < 0.05, ** = p < 0.01, *** = p < 0.001; Kruskal-Wallis test with Dunn’s post hoc and Holm-Bonferroni correction).

To further dissect these changes in *alkB* promoter activity, we analysed the top 10% most fluorescent cells and compared their fluorescence relative to baseline levels **(Fig 4B)**. In both experiments, median relative fluorescence of this subpopulation was higher than baseline at day two (one-sample t-test, experiment 1: *t*(7) = 8.36, *p* = 6.9 × 10^-5^; experiment 2: *t*(7) = 3.03, *p* = 0.0191) and day seven (experiment 1: *t*(7) = 6.56, *p* = 3.2 × 10^-4^; experiment 2: *t*(7) = 4.84, *p* = 1.9 × 10^-3^). Relative fluorescence also varied between time points (Kruskal-Wallis, *H*(2) = 2, *p* = 0.368), with significantly higher fluorescence at day two compared to time zero (Dunn’s test, *p*_*Exp1*_ = 0.00612; *p*_*Exp2*_ = 0.0197). Additionally, in the second experiment, the median relative fluorescence increased twofold on day seven compared to time zero (Dunn’s test, *p =* 0.00235). This analysis confirmed that while median fluorescence of the total population remained low, a subset of highly fluorescent cells maintained or even increased their fluorescence relative to baseline levels.

Overall, these results showed that PFF2_P*alkB*-GFP*_ single-cell fluorescence heterogeneity increased in the phyllosphere over time. While median fluorescence was low, subpopulations of highly fluorescent cells persisted, suggesting that *alkB* promoter activation is differentially regulated within a population.

## Discussion

The phyllosphere represents harsh conditions for microbes to live on, including the low and heterogeneous availability of easily-accessible carbon sources like sugars (Leveau and Lindow 2001a; Remus-Emsermann et al. 2011). Another potential carbon source for bacteria could be the plant cuticle itself, which consists of alkanes. PFF2 encodes for the *alkB* gene, which is associated with alkane degradation (Vomberg and Klinner 2000). Additionally, it has been shown that PFF2 is able to utilise diesel better compared to its *alkB* gene knockout mutant indicating that *alkB* is actively expressed (S. Oso et al. 2021). Therefore, we tested the hypothesis that PFF2 upregulates *alkB* on the waxy cuticle of plants, potentially to utilise the alkanes in the waxes covering the cuticles of plants (Zeisler-Diehl, Barthlott, and Schreiber 2020). To test this, we developed a bioreporter in the PFF2 background (PFF2_P*alkB*-GFP*_) that expresses the green fluorescent mClover3 protein under the control of the PFF2 *alkB* promoter and is destabilised by adding an alanin-alanin-valin peptide (Leveau and Lindow 2001b). As a result, the fluorescent protein is degraded shortly after its maturation, allowing the bioreporter to reflect the current promoter activity, compared to the usually high stability of fluorescent proteins in bacteria.

As previously shown for its wild type, PFF2_P*alkB*-GFP*_ can grow on diesel as its sole carbon source (S. Oso et al. 2021). However, from day 18 onwards, the OD_600nm_ of PFF2_P*alkB*-GFP*_ decreased despite the presence of diesel droplets. This is likely a result of nutrient depletion, as only a fraction of diesel is bioavailable for bacterial utilisation (Xu et al. 2018). During growth on diesel, the *alkB* promoter was upregulated from day one onwards compared to growth on BHB minimal medium without a carbon source. By day 28, similarly to the observed optical density, the promoter activity declined. Notably, the *alkB* promoter activity was not uniformly distributed across the population. Instead, a large proportion of the population displayed minimal expression of *alkB*, with fluorescence distributions consistently right skewed across all conditions. Over time, 46–90% of diesel-grown cells exhibited fluorescence values below 10% of the maximum measurable intensity, further supporting non-uniform *alkB* promoter activity. This heterogeneity indicates a potential bet-hedging strategy, where subpopulations adopt distinct metabolic states, some of which actively engage in *alkB* expression and presumably alkane degradation, while others may rely on alternative metabolic pathways. The presence of low-expressing cells across all conditions indicates that a subset of the population may persist in a dormant or alternative metabolic state, possibly scavenging metabolites released by high-expressing cells during growth on alkanes (Kamrad et al. 2023). In LB and BHB without diesel, most PFF2_P*alkB*-GFP*_ cells exhibited a low *alkB* promoter activity. However, a subset of cells displayed higher-than-expected fluorescence, suggesting that *alkB* expression is not entirely repressed under non-alkane conditions. This could be due to stochastic gene expression, presence of trace alkanes in the culture flasks, or alternative regulatory mechanisms influencing *alkB* activation.

In planta, approximately 10–25% of PFF2_P*alkB*-GFP*_ cells actively expressed GFP* at two- and seven-days post-inoculation, while the remaining showed low *alkB* promoter activity. This indicates that most of the population focused on alternative metabolic pathways for growth, such as sugar metabolism (Leveau and Lindow 2001a), or remained metabolically inactive. Unlike bacteria reporting on the activity of the *fruR* promoter, which indicated a rapid depletion of fructose and sucrose within the first 24 hours (Leveau and Lindow 2001a), the *alkB* promoter was found to be active in a subset of the population for seven days. This indicates that the available alkane concentration fluctuates less than sugar photosynthates, which leach from the inside of the leaf to the outside (Tukey and Mecklenburg 1964).

Over the timespan of seven days, the fluorescence of PFF2_P*alkB*-GFP*_ were increasing. An explanation for this could be that when the bacteria were first inoculated onto the plant, there was a high availability of sugars on the plant surface and, due to carbon catabolite repression, these primary carbon sources were utilised before alkanes. These sugars, including sucrose, fructose, and glucose (Mercier and Lindow 2000), are photosynthates that become less available over time as they are consumed by bacteria (Leveau and Lindow 2001a). To ensure survival, bacteria may need to rely on another carbon source. Therefore, they could start degrading alkanes, leading to the observed higher P*alkB* activity. It has been shown that some leaf areas feature hotspots of diffusion, leading to higher availability of photosynthates (Remus-Emsermann et al. 2011, 2012; Schlegel, Schönherr, and Schreiber 2005). Consequently, bacterial populations are larger at those sites. This could explain the large subpopulation of cells that barely activated *P*alkB compared to others which did not have access to sugar photosynthates but could access alkanes.

Throughout the experiment, fluorescence intensity varied widely among cells. This heterogeneity could be attributed to the high spatial and nutrient heterogeneity of leaf surfaces (Remus-Emsermann and Schlechter 2018). In some areas such as the bases of trichomes, above veins, and in epidermal cell grooves, there are higher concentrations of sugars (Schlechter, Miebach, and Remus-Emsermann 2019). This initially reduced the need for alkane degradation in bacteria to survive. Also, accessibility of alkanes in the cuticle may differ across the leaf surface, further contributing to the heterogeneous *alkB* expression in PFF2. Further experiments using fluorescence *in situ* microscopy on leaves, instead of washed off bacteria, could provide more information about whether highly fluorescent cells are spatially arranged in relation to conspecific cells and on the leaf environment.

## Conclusion

Previous studies have shown that bacteria carrying *alkB* genes are prevalent on leaf surfaces. However, the role of *alkB* in utilising alkanes and other aliphatic components from leaf cuticles remains unclear. Here, we have provided further evidence that subpopulations of bacteria on leaves can activate *alkB* expression, indicating its role in resource utilisation when other resources may be scarce. However, direct functional evidence of cuticular wax degradation by phyllosphere bacteria is still missing. Our results indicate that bacteria may access cuticular waxes from the leaf, but further studies need to confirm this. Future experiments should focus on this in further detail, e.g. by measuring the incorporation of radiolabelled cuticular waxes during bacterial colonisation of leaves. Additionally, the activity of other proxy genes for aliphatic compounds, such as *almA*, should be investigated to have a broader understanding of cuticular wax utilisation and bacterial adaptations to the phyllosphere.

## Author Contributions

MRE and RS conceived and planned the study. LV, EK, MO, RS performed experiments. RS and LV analysed data. RS, LV, and MRE wrote the manuscript. EK and MO gave feedback on and revised the manuscript.

## Acknowledgments

The authors thank Sandra Hirsch for her valuable technical assistance and for maintaining the day-to-day operations of the laboratory.

## Conflict of Interest

The authors declare no conflict of interests.

## Code and data availability

All images and data supporting this study are available in Zenodo (Voß et al. 2025). The R scripts used for data analysis are accessible in the following GitHub repository: https://github.com/relab-fuberlin/alkanebioreporter.

## Notes

https://github.com/relab-fuberlin/alkanebioreporter

https://zenodo.org/records/15014571

### Competing Interest Statement

The authors have declared no competing interest.

