## Supplemental Material for "The alkane 1-monooxigenase gene *alkB* of *Pseudomonas* sp. FF2 is upregulated during colonisation of *Arabidopsis thaliana* leaves"

Supplemental Methods

#### **Construction of pPalkB-GFP***

The plasmid pFru97-PalkB-mClover3-[AAV] (pPalkB-GFP*) was constructed via isothermal assembly (Gibson et al. 2009). A 300-bp region upstream of the *alkB* gene (P*alkB*) from PFF2 was amplified using primers PFF2_alkB-for (gccaggaattggggatcggaAATCGTGCCATTTCATTCGCG) and PFF2_alkB-rev (ccttgctcatGGCAACCTCCGAAGGC) using touchdown PCR. The green fluorescent protein coding *mClover3* gene was amplified from pMRE137 (Bajar et al. 2016; Schlechter et al. 2018), using primers mClover3_PFF2_fw (CCGCAATGAGGACCACGATTATTatgagcaagggcgaggag) and XFPAAV_RV. The primer XFP-AAV_RV (gtccaagctcagctaaTTACACGGCGGCcttgtacagctcgtcc) adds a short degradation peptide signal (alanin-alanin-valin) at the C-terminus of mClover3, resulting in destabilisation of the fluorescent protein (Leveau and Lindow 2001b). The P*alkB* fragment was fused to the promoterless *mClover3* gene by overlap extension PCR using primers PFF2_alkB-for and XFP-AAV_RV, resulting in a 1-kb fragment. The plasmid backbone pFru97 (Tecon and Leveau 2012) was linearised with HindIII (New England Biolabs) and the PalkB-mClover3-[AAV] construct was inserted into the plasmid using isothermal assembly (Gibson et al. 2009). The assembled pPalkB-GFP* plasmid was then transformed into NEB Turbo competent *E. coli* cells (New England Biolabs) according to the manufacturer’s recommendations and were selected on LBA supplemented with Km. The plasmid was verified by Sanger sequencing (GENEWIZ Germany GmbH) and then transformed into chemically competent *E. coli* ST18 before plating on LBA supplemented with Km + 5-aminolevulinic acid (5-ALA, 50 mg/ml) to select for transformants (Thoma and Schobert 2009).

#### **Conjugation of pPalkB-GFP* into PFF2**

To transfer pPalkB-GFP* into PFF2, biparental mating using *E. coli* ST18(pPalkB-GFP*) as donor strain was performed. Biparental mating was conducted as described in Schlechter *et al.* (2019). Briefly, donor and recipient strains were mixed in a 1:1 ratio based on optical density at 600 nm (OD_600nm_), drop spotted onto LBA + 5-ALA, and incubated overnight at 30°C. Then, the bacterial mix was harvested and resuspended in phosphate buffered saline (1×PBS; 8 g L^-1^ NaCl, 0.2 g L^-1^ KCl, 1.44 g L^-1^ Na_2_HPO_4_, 0.24 g L^-1^ KH_2_PO_4_; pH 7.2). Selection for pPalkB-GFP* was conducted on LBA + Km at 30°C. Media devoid of 5-ALA counterselects against the donor *E. coli* ST18 strain. Positive colonies were screened via PCR for *E. coli* contamination by targeting the conserved β-D-glucuronidase enzyme-coding gene in *E. coli* with primers uidA_FW (AACAGGTGGTTGCAACTGGA) and uidA_RV (TTGCTGAGTTTCCCCGTTGA). To test for the presence of plasmid pPalkB-GFP*, REV_pFru_ins (ATAAACTGCCAGGAATTGGGG) and FWD_pFru_ins (CAACAGGAGTCCAAGCTCAG) were used, as these primers target a fragment of the plasmid backbone sequence (Schlechter et al., 2018). A positive PFF2(pPalkB-GFP*) transconjugant was selected for further analysis from here onwards referred to as PFF2_P_*_alkB_*_-GFP*_.

### **Growth of PFF2_P_*_alkB-GFP*_ in planta* and activity of P*alkB***

To determine *alkB* promoter activity during colonisation of *A. thaliana* leaves, PFF2_P_*_alkB_*_-GFP*_ was inoculated onto axenically grown *A. thaliana* as described previously (Miebach et al. 2020). *A. thaliana* Col-0 seeds were surface-sterilised in 70% v/v ethanol for two minutes. Then, ethanol was removed, and seeds were treated with 50% v/v bleach solution (NaOCl, 2.47% w/w) with 0.02% v/v Tween-20 for seven minutes while mixing the seeds regularly. After discarding the bleach solution, the seeds were washed three times with sterile distilled water and stratified at 4°C in the dark for two days. Afterwards, the seeds were sown onto ½ strength Murashige and Skoog (MS with vitamins, pH 5.8, Duchefa) 1.0% w/v plant agar (Duchefa) plates with each seed placed on a trimmed sterile pipette tip filled with solid MS agar. These plates were then sealed with micropore tape (3M Deutschland GmbH) and transferred into a plant growth cabinet (poly klima GmbH). Seeds were germinated at 21 °C in an 11/13 h photoperiod, with a light intensity of ~120 μE m^-2^ s^-1^, and a relative humidity of 80%. Eight days after sowing, the seedlings were transferred aseptically into autoclaved Magenta GA-7 (Merck) plant tissue culture boxes containing 90 g of zeolite (Zeolith-100, Steinlando) and 45 ml of sterile ¾ MS. Four seedlings were placed per box. The boxes were then further incubated in the plant growth cabinet for three weeks prior to inoculation.

The inoculum was prepared by growing an overnight culture of PFF2_P_*_alkB_*_-GFP*_ (3 ml LB + Km). Then, a fresh culture was prepared by inoculating 50 ml of LB + Km with 500 μl of overnight culture. The culture was then incubated at 30°C at 300 r.p.m. until entering the log-phase (approximately three hours). Three millilitres of culture was then washed by centrifugation (5 min at 5,000 × *g*) and resuspended in the same amount of sterile 1×PBS to an OD_600nm_ of 0.05. The suspension was then sprayed onto six magenta boxes (200 µL per box) using an airbrush gun (Ultra airbrush, Harder & Steenbeck GmbH & Co. KG; compressor, Sparmax TC-620X airbrush compressor). The inoculated boxes were then placed into the plant growth cabinet under the same controlled conditions.

Whole *A. thaliana* rosettes were sampled on day zero, two, and seven after inoculation, with eight biological replicates sampled each time. After cutting off the roots, plants were placed into a 15-ml tube, and plant fresh plant weight was determined before 1 ml of sterile 1 × PBS was added. Samples were vortexed for 10 sec, sonicated for 5 minutes at 75% intensity in a sonication bath (Emmi-12HC, EMAG AG), and then vortexed for an additional 15 sec. An aliquot of the leaf wash was used to prepare a tenfold dilution series and to determine colony forming units (CFU) on LBA + Km. The remaining suspension was used for microscopy.

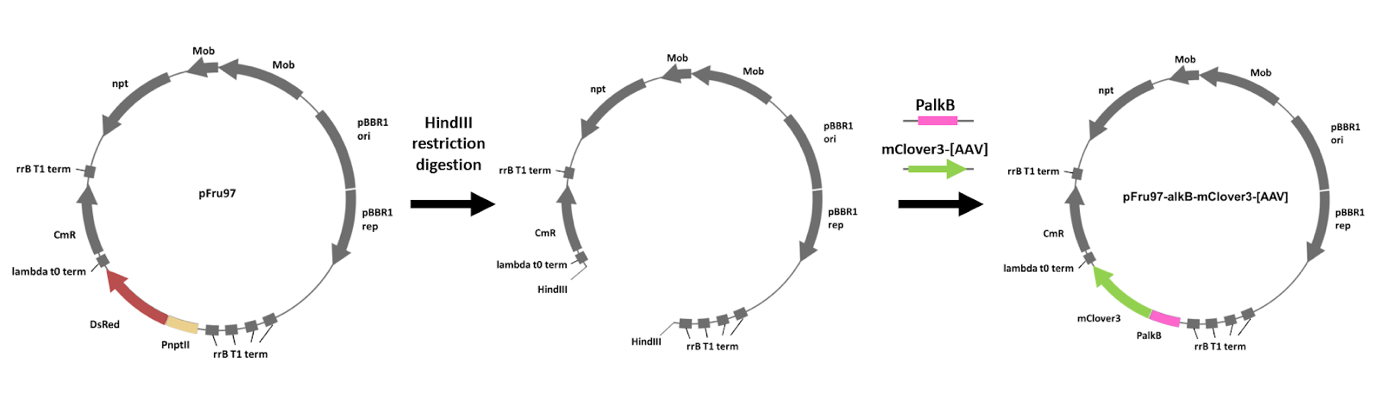

**Figure S1. Plasmid construction.** The *alkB* promoter of *Pseudomonas* sp. FF2 and green fluorescence protein gene mClover3-[AAV] were amplified and inserted into the backbone of the HindIII-digested pFru97 expression plasmid through Gibson assembly, resulting in plasmid pFru97-alkB-mClover-[AAV] (pPalkB-GFP*).

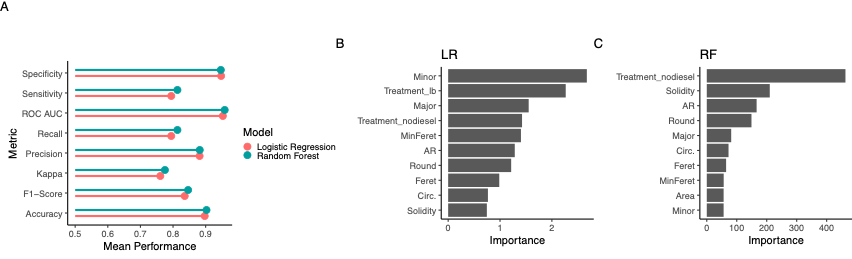

**Figure S2. Comparison of supervised machine learning models for *in vitro* single-cell data analysis.** (A) Performance metrics of logistic regression (LR) and random forest (RF) models for cell classification of PFF2 cells grown in vitro (BHB or LB medium). Feature importance of (B) LR and (C) RF models. Features: Treatment_X1 (BHB medium); Treatment_X2 (LB medium); Minor (minor axis length); Major (major axis length); MinFeret (minimum feret diameter); Feret (maximum feret diameter); AR (aspect ratio); Round (roundness); Circ. (circularity); Solidity (area/convex hull area ratio); Area (total number of pixels within a particle).

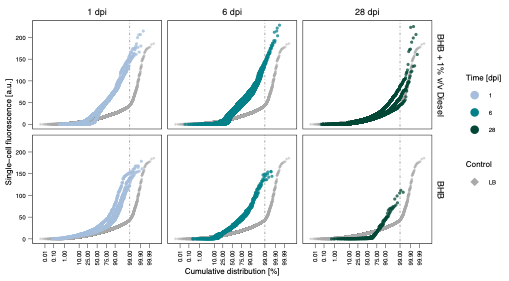

**Figure S3. Distribution of PFF2_PalkB-GFP*_ single-cell GFP fluorescence *in vitro*.** Normal probability plot of cell fluorescence of PFF2_PalkB-GFP*_ grown in BHB supplemented with or without diesel over time (1, 6, and 28 days) or LB. Vertical line indicates the 99% of cells. Single-cell fluorescence of PFF2_PalkB-GFP*_ grown in LB fluorescence was included for comparison (gray diamond).

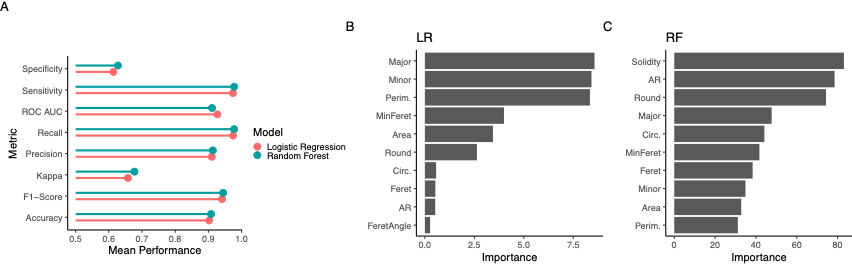

**Figure S4. Comparison of supervised machine learning models for *in planta* single-cell data analysis.** (A) Performance metrics of logistic regression (LR) and random forest (RF) models for cell classification of PFF2 cells grown in planta (A. thaliana phyllosphere). Feature importance of (B) LR and (C) RF models. Features: Minor (minor axis length); Major (major axis length); Perim. (perimeter); MinFeret (minimum feret diameter); Feret (maximum feret diameter); FeretAngle (angle of the maximum feret diameter); AR (aspect ratio); Round (roundness); Circ. (circularity); Solidity (area/convex hull area ratio); Area (total number of pixels within a particle).

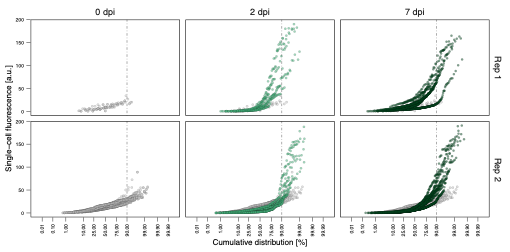

**Figure S5. Distribution of PFF2_PalkB-GFP*_ single-cell GFP fluorescence in the *Arabidopsis phyllosphere*.** Normal probability plot of PFF2_PalkB-GFP*_ single-cell GFP fluorescence at different sampling points (0-, 2-, and 7-days post-inoculation). Each point represents the fluorescence intensity of an individual cell. Fluorescence at time zero is included in every plot as a reference (gray).
